# Subcortical Shape Alterations in Major Depressive Disorder: Findings from the ENIGMA Major Depressive Disorder Working Group

**DOI:** 10.1101/534370

**Authors:** Tiffany C. Ho, Boris Gutman, Elena Pozzi, Hans J. Grabe, Norbert Hosten, Katharina Wittfeld, Henry Völzke, Bernhard Baune, Udo Dannlowski, Katharina Förster, Dominik Grotegerd, Ronny Redlich, Andreas Jansen, Tilo Kircher, Axel Krug, Susanne Meinert, Igor Nenadic, Nils Opel, Richard Dinga, Dick J. Veltman, Knut Schnell, Ilya Veer, Henrik Walter, Ian H. Gotlib, Matthew D. Sacchet, André Aleman, Nynke A. Groenewold, Dan J. Stein, Meng Li, Martin Walter, Neda Jahanshad, Paul M. Thompson, Philipp G. Sämann, Lianne Schmaal

## Abstract

Alterations in regional subcortical brain volumes have been widely investigated as part of the efforts of an international consortium, ENIGMA, to determine reliable structural brain signatures for Major Depressive Disorder (MDD). Given that subcortical structures are comprised of distinct subfields, we sought to build significantly from prior work to precisely map localized MDD-related differences in subcortical regions using shape analysis. In this meta-analysis of subcortical shape from the ENIGMA-MDD working group, we compared 1,781 patients with MDD and 2,953 healthy controls (CTL) on individual measures of shape metrics (thickness and surface area) on the surface of seven bilateral subcortical structures: nucleus accumbens, amygdala, caudate, hippocampus, pallidum, putamen, and thalamus. Harmonized data processing and statistical analyses were conducted locally at each site, and findings were aggregated by meta-analysis. Relative to CTL, patients with MDD had lower surface area in the subiculum of the hippocampus, the basolateral amygdala, and the nucleus accumbens shell. Relative to CTL, patients with adolescent-onset MDD (≤ 21 years) had lower thickness and surface area of the subiculum of the hippocampus and the basolateral amygdala. Relative to first-episode MDD, recurrent MDD patients had lower thickness and surface area in the CA1 of the hippocampus and the basolateral amygdala. Our results suggest that previously reported MDD-associated volumetric differences may be localized to specific subfields of these structures that have been shown to be sensitive to the effects of stress, with important implications for mapping treatments to patients based on specific neural targets and key clinical features.

## Introduction

Major Depressive Disorder (MDD) is one the leading causes of disability worldwide, with relatively high rates of lifetime prevalence and recurrence (World Health 2017). MDD is often triggered by stressful experiences and is commonly associated with various affective symptoms (e.g., abnormalities in emotion regulation, reduced motivation in the face of positive incentives, sustained experiences of negative affect (Davidson et al 2002; Woody and Gibb 2015), as well as with cognitive deficits (e.g., attention, learning, working memory, processing speed, motor functioning (McIntyre et al 2013)). Several subcortical regions – particularly the hippocampus, amygdala, and structures of the striatum – through their connections with one another and with cortical structures, are important for supporting a number of these cognitive and affective processes that are disturbed in MDD (Davidson et al., 2002). In a recent multi-site effort, we examined morphological alterations at the level of subcortical gray matter volumes in MDD (Schmaal et al 2016), and found lower total hippocampal volumes, mainly driven by patients with recurrent episodes and by patients with a relatively early age of onset (i.e., prior to age 21). Despite the large study sample size and homogeneous analysis protocols, no statistically significant group differences emerged for any of the other subcortical structures. It is possible, however, that such aggregate measures of volume are either insensitive to local volumetric effects or that they obscure heterogeneous local effects by averaging out more complex shape effects. In this respect, the analysis of shape parameters may represent a more sensitive approach. Beyond this, local variations in shape measures are highly heritable, and specific genetic variants may affect only localized regions of the structure (Hibar et al 2015; Roshchupkin et al 2016), but also to provide insight into the anatomical relation with important clinical variables, such as illness onset and recurrence, as well as the ability to detect fine-grained changes that may be particularly helpful in the context of monitoring intervention targets with more specificity. Indeed, all of the subcortical structures that have been implicated in MDD contain functionally distinct subregions (Roddy et al 2018). For example, the hippocampus is not a singular unit, but consists of several subregions, including the cornu ammonis subfields (CA) 1-4, dentate gyrus (DG), and the subiculum (SUB). While the more dorsal regions are involved in memory formation and spatial cognition, the ventral regions subserve affective processing (Fanselow and Dong 2010). Most hippocampal subregions are sensitive to effects of glucocorticoids and psychosocial stress (Teicher et al 2012; Wang et al 2013), with reported reductions in individuals with MDD compared to healthy controls, particularly in CA1, CA3, DG, and SUB (Bearden et al 2009; Boldrini et al 2010; Han et al 2016; Treadway et al 2015). Importantly, there is evidence from both functional and structural imaging that these stress-sensitive subfields of the hippocampus are especially vulnerable to environmental factors that influence their development (Teicher et al 2012). In this context, early life stress may explain prior reports of reduced hippocampal volume in MDD (Frodl et al 2016; Schmaal et al 2016) by influencing hippocampal development to increase risk for MDD.

Similarly, the amygdala contains functionally distinct subregions. Based on anatomical tracing studies and cellular architecture, two primary nuclei subpopulations in the amygdala have been identified: the basolateral and centromedial amygdala (BLA, CMA) (Mosher et al 2010; Sah et al 2003). The CMA, with its reciprocal connections with the basal ganglia, midbrain, and brain stem, appears to be involved in allocating attention and generating the appropriate autonomic responses to environmental stimuli. In contrast, the BLA, with its connections to the CMA and with extensive cortical regions, is primarily involved in evaluating the emotional content of sensory inputs and plays a major role in threat responses (Mosher et al 2010; Terburg et al 2018). Although some post mortem evidence suggests that MDD selectively affects the receptors and number of nuclei in the BLA (Karolewicz et al 2009; Rubinow et al 2016), it is not clear if MDD is associated with larger or smaller BLA volume. Several confounding variables related to disease burden (e.g., age of illness onset, recurrence, medication usage) likely contribute to these discrepant results (Hamilton et al 2008; Kronenberg et al 2009; Rubinow et al 2016; van Eijndhoven et al 2009). Another possibility is that associations of MDD with amygdala volumes are specific to certain subdivisions and are not easily detected when the total amygdala volume is used (Schmaal et al 2016).

Finally, the striatum is comprised of the dorsal striatum (caudate, putamen) and ventral striatum (nucleus accumbens; NAcc). Cytoarchitectural, histological, and functional imaging evidence suggests that the caudate head is more strongly related to affective and cognitive processes, whereas the caudate body/tail is more centrally involved with processing sensory inputs to shape perception and action (Robinson et al 2011). Similarly, the ventral putamen has a more prominent role in processing emotions in a motivational context, whereas the dorsal putamen, along with other nuclei in the basal ganglia, is primarily responsible for motor learning (Haber 2016; Haber and Knutson 2010). The NAcc – situated adjacent to the medial and ventral parts of the caudate and putamen – contains a shell (NAcc-s) and a core. The nuclei of the NAcc-s are structurally distinct from those in the core and have different functional roles (Bertran-Gonzalez et al 2008; Heimer et al 1991). While neurons in the core project to the pallidum, substantia nigra, and other motor areas, NAcc neurons project to the extended amygdala and ventral tegmental area that are involved in processing pleasure signals, motivational salience, and reward-based reinforcement (Richard et al 2013). Aversive stressful events may lead to striatal dopamine release, underscoring a role of the striatum in stress-related diseases including MDD (Richard et al 2013). Structural alterations in these striatal subregions have also been associated with motivation- and reward-related disturbances – ranging from blunted responses to positive reinforcers and biases towards negative feedback – in patients with MDD (Francis et al 2015; Martin-Soelch 2009; Pizzagalli et al 2009).

MDD is most likely characterized by specific associations with functionally distinct subregions within the hippocampus, amygdala, striatum, and other subcortical structures (e.g., thalamus). It is important to note, however, that characterizing local patterns in subcortical surfaces has traditionally been challenging due to the lack of identifiable surface landmarks that are more common in cortical surfaces (e.g., deep sulcal patterns). Further, relying on smaller samples from individual sites may not provide sufficient statistical power to detect subtle effects in subcortical shape or to reliably overcome variation due to methodological (e.g., preprocessing pipeline, scanner idiosyncrasies) or clinical factors (e.g., age of illness onset, recurrence, and medication exposure). In this context, the lack of detectable volumetric differences in the amygdala, caudate, putamen, and NAcc between MDD and CTL in our previous meta-analytic study may be due to the fact that we did not use shape analyses to examine these important subdivisions (Schmaal et al 2016).

To address these knowledge gaps, we conducted a multi-site meta-analytic investigation to test whether MDD patients, and whether specific subgroups of MDD based on important clinical characteristics, show differences from controls in subcortical shape. Specifically, we applied meta-analytic models on effect sizes generated from 10 study cohorts from 6 different countries participating in the MDD Working Group of the international ENIGMA consortium. Each study site applied a well-validated harmonized preprocessing pipeline and conducted statistical models on high-resolution T1-weighted MRIs, yielding site-level summary statistics of volume and shape for seven bilateral subcortical regions from 1,781 patients diagnosed with MDD and 2,953 healthy controls (CTL).

Guided by findings from our prior meta-analysis in which we reported that the most robust difference between individuals with MDD and CTL was smaller hippocampal volume (Schmaal et al., 2015), and from recent work indicating that CA1, CA3, DG, and SUB are associated with exposure to aversive stressful experiences (Teicher et al 2012) and MDD (Han et al 2016; Treadway et al 2015), we hypothesized that patients with MDD would exhibit reductions in these hippocampal subregions. Given previously documented effects of age of illness onset and recurrence of illness on subcortical volumes (primarily the amygdala and hippocampus; Hamilton et al 2008; Schmaal et al 2016), we also sought to stratify groups according to these clinical characteristics: early (prior to age 21) versus later (after age 21) onset MDD and first-episode versus recurrent-episode. We also report additional exploratory analyses of medicated and non-medicated patients (each compared separately to CTL) and dimensional associations between subcortical shape and depression severity (clinician-rated as well as self-reported) among patients with MDD.

## Materials and Methods

### Samples

Ten participating sites in the MDD Working Group of ENIGMA consortium (Schmaal et al 2017; Schmaal et al 2016; Thompson et al 2014) applied harmonized preprocessing and statistical models on structural T1-weighted MRIs, yielding site-level summary statistics of subcortical volume and shape from a total of 4,734 participants (1,781 patients with MDD and 2,953 CTL). Detailed demographics, clinical characteristics, and exclusionary criteria for study enrollment for each site are presented in supplemental Table S1. All participating sites obtained approval from their respective local institutional review boards and ethics committees. All study participants provided written consent at their local site.

### Clinical variables of interest

We selected specific clinical variables of interest based on prior work demonstrating their effects on aggregate subcortical volumes in MDD (Hamilton et al 2008; Schmaal et al 2016). These variables included: age of illness onset, number of episodes, and the severity of depressive symptoms at the time of scan as measured by the clinician-rated 17-item Hamilton Depression Rating Scale (HDRS-17; Hamilton 1960) or the 21-item self-report Beck Depression Inventory (BDI; Beck et al 1961). Consistent with our prior work, we considered participants with earlier or adolescent onset (EO) to be those who developed first episodes at or before age 21, and participants with later or adult onset (LO) to be those who developed first episodes after age 21 (Schmaal et al 2016). We defined recurrent-episode MDD (RECUR) to be those who experienced more than one major depressive episode (Schmaal et al 2016). As a supplemental analysis (because the majority of sites did not include detailed information on lifetime medication usage, dosage, or adherence), we also compared MDD groups on the basis of antidepressant medication usage at the time of scan.

### Image processing and analysis

All participating sites collected anatomical T1-weighted MRI brain scans locally at each site, where they were analyzed using the fully-automated and validated segmentation software FreeSurfer version 5.3 (Fischl 2002), with the exception of 3 sites which used version 5.0 or 5.1 (see Table S1). A subset of these subcortical measures have been previously published (Frodl et al 2016; Renteria et al 2017; Schmaal et al 2016); however, none of these prior meta-analyses from the MDD Working Group of ENIGMA conducted shape analyses. Image acquisition parameters and software descriptions for each sample are presented in supplemental Table S1. The 7 bilateral subcortical segmentations were: the nucleus accumbens, amygdala, caudate, hippocampus, pallidum, putamen, and thalamus (as well as lateral ventricles and total intracranial volume, ICV). All segmentations were visually inspected for accuracy following standardized protocols (http://enigma.ini.usc.edu/protocols/imaging-protocols/).

We analyzed shape using the ENIGMA-Shape protocol (http://enigma.usc.edu/ongoing/enigma-shape-analysis/), for which test-retest reliability has been previously validated (Hibar et al., 2017). Briefly, shapes were extracted using the FreeSurfer 5.3 parcellation, followed by a topological correction and mild smoothing based on the topology-preserving level set algorithm (Gutman et al 2015). As in prior work, after registering shapes to standardized templates, we then defined two vertex-wise measures of space morphometry which facilitated comparisons of subcortical shape: *radial distance*, as derived from the medial model (Gutman et al 2015; Gutman et al 2012), which yields a measure of “shape thickness,” and the *Jacobian determinant*, as derived from tensor based morphometry (TBM; Gutman et al 2015; Wang et al 2011), which yields a metric of localized tissue reduction or enlargement of surface area (relative to the respective template shape). Because the Jacobian represents the ratio of the area in the individual shape relative to the area in the template at the corresponding vertex – and because this is not Gaussian in distribution – we used the logarithm of the Jacobian in all analyses examining shape surface area, as this tends to be closer to Gaussian. A useful feature of the ENIGMA-Shape pipeline is that results are based on bilateral shape measures (i.e., templates for corresponding left and right regions are vertex-wise registered after reflecting one of them, and summed vertex-wise). Importantly, our registration algorithm provides a unique and stable matching between datasets, allowing us to efficiently meta-analyze the effects of MDD across datasets (as in Roshchupkin et al 2016). See “Image processing and analysis” under the Supplemental Information for more details on the pipeline for subcortical shape analysis and on quality control procedures.

Each of the 10 study sites applied the subcortical shape pipeline and ran *a priori* statistical models (for details, see “Statistical framework for meta-analyses”, below) that were guided by discussions with ENIGMA-MDD members and previous work (Schmaal et al 2017; Schmaal et al 2016) to generate summary statistics for inclusion in our meta-analyses.

### Site-specific statistical models

To harmonize analyses across sites, a set of standardized scripts to compute mass univariate statistics was distributed to all participating sites via the ENIGMA-Git page (https://github.com/ENIGMA-git/ENIGMA/tree/master/WorkingGroups). Each study site performed mass univariate (per-vertex, per-measure) analysis for all the linear models proposed in the present study (see Table 1). Specifically, for our primary statistical models of interest, subcortical shape measures of thickness (radial distance) and surface area (log of the Jacobian determinant) were the outcome variables, and a binary group indicator variable (e.g., 0=CTL, 1=MDD) was the predictor of interest, with age, sex (as a factor), and total ICV as covariates. Secondary analyses included the following planned comparisons: early onset MDD (EO) vs. CTL; later onset MDD (LO) vs. CTL; EO vs. LO; recurrent episode MDD (RECUR) vs. CTL; first episode MDD (FIRST) vs. CTL; RECUR vs. FIRST. We also tested for associations with HDRS-17 and BDI scores (separately) within the MDD group only. Our exploratory analyses included comparing groups based on antidepressant usage at the time of scan (MED vs. CTL, and NON vs. CTL), as well as testing whether sex and age significantly interacted with diagnostic group in the primary analysis (MDD vs. CTL) to explain variation in subcortical shape measurements.

**Table 1.**
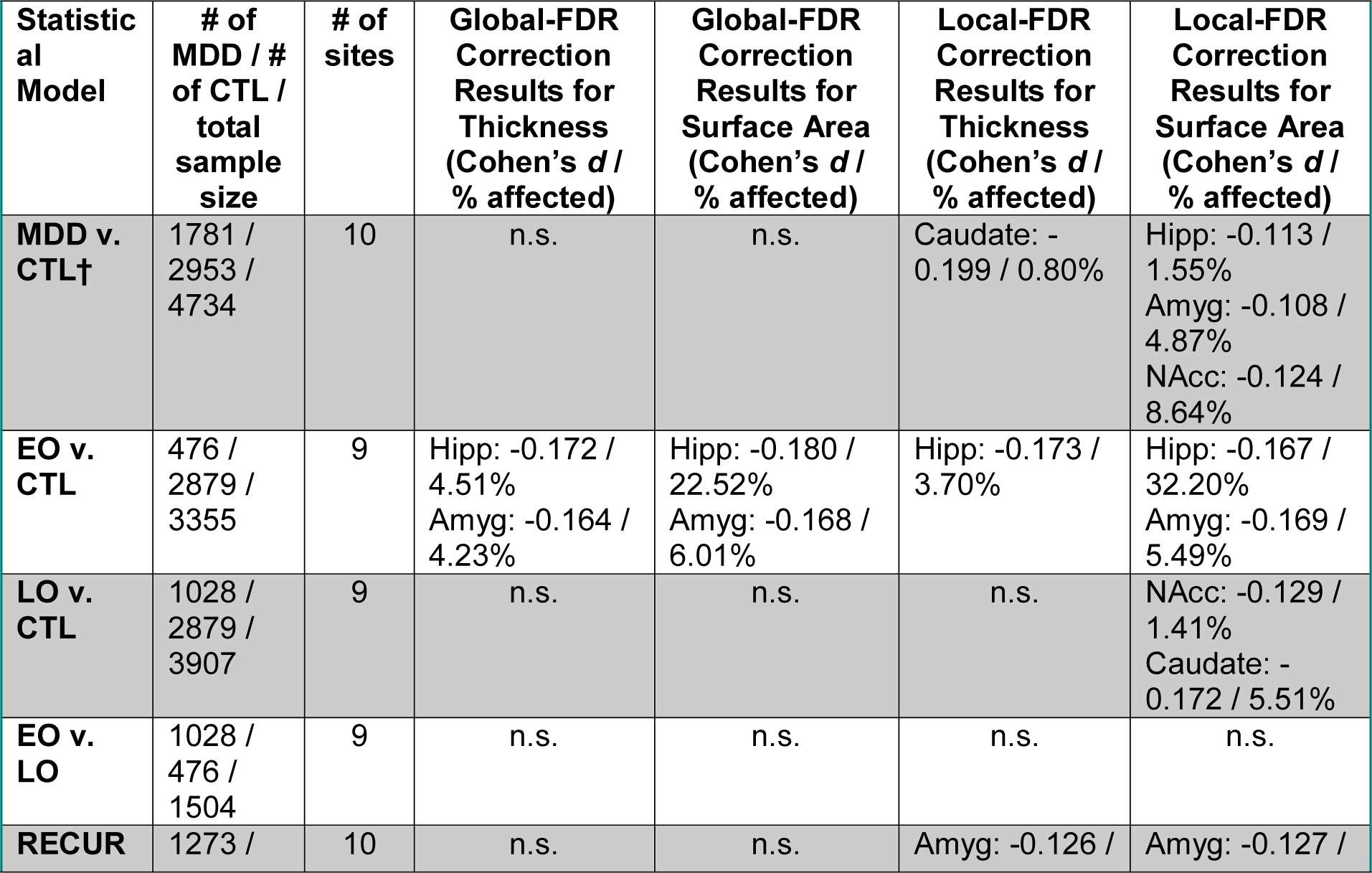
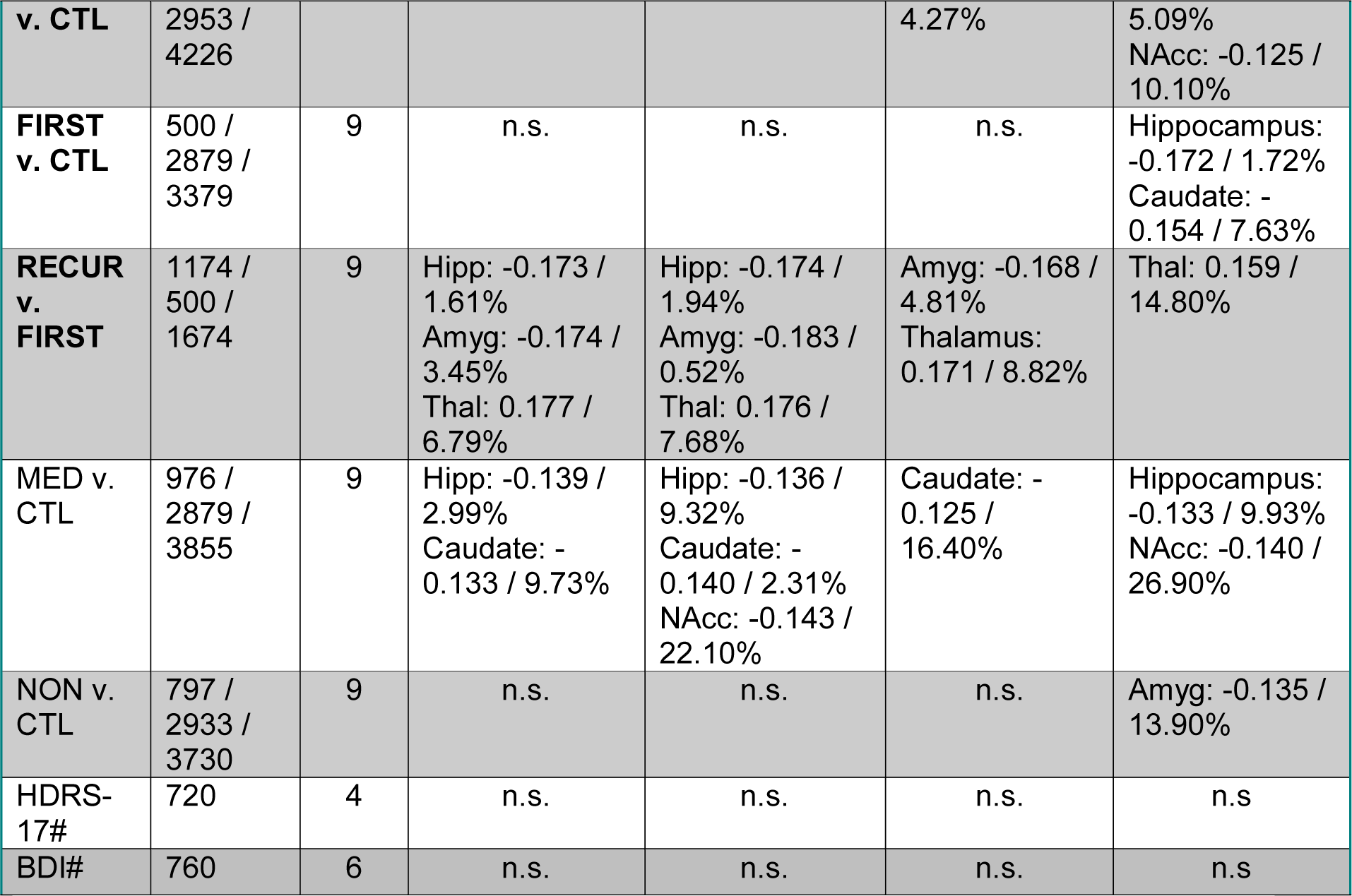
List of primary (bolded) and supplemental statistical analyses. All site-specific analyses included age, sex (as a factor), and intracranial volume (ICV) as covariates and all meta-analytic models pooled each sample’s effect sizes (i.e., *d* or *r*) using an inverse variance-weighted random effects model. For more information on each study site, please see Table S1. Thickness is measured by radial distance and surface area is measured using tensor-based morphometry. See Figures 1-2 for more details on results from the primary analyses surviving global-FDR correction, Figures 3 and S1-S4 for more details of results from the primary analyses surviving local-FDR correction, and Figures S5-S7 for results on the supplemental analyses. MDD=Major Depressive Disorder; CTL=healthy controls; EO=early-onset MDD (≤ 21 years old); LO=later-onset MDD (> 22 years old); FIRST=first-episode MDD; RECUR=recurrent-episode MDD; MED=medicated at time of scan; NON=not medicated at time of scan; HDRS-17=Hamilton Depression Rating Scale (17 items); BDI=Beck’s Depression Inventory; n.s.=no significant effects; †interactions between age and sex (separately) were also tested; # indicates dimensional analyses conducted within MDD only.

### Meta-analytic framework and correction for multiple comparisons

The resulting group-level maps of effect sizes (i.e., Cohen’s *d* for the group comparisons and Pearson’s *r* for the dimensional analyses), regression parameters, and confidence intervals, as well as basic site information, were aggregated for mass univariate meta-analysis. As performed in Schmaal et al 2017; Schmaal et al 2016, we conducted meta-analyses which pooled each site’s effect sizes, for each region, using an inverse variance-weighted random-effects model as implemented in the R package *metafor* (version 1.9-1) and fit with REML (https://cran.r-project.org/). One advantage of random effects models is that they allow effect sizes to vary across studies due to study-specific differences (e.g., mean age); random effects models therefore weight within-study as well as between-study variance in the pooled effect size estimates to mitigate bias or undue influence from the largest samples in the meta-analysis (Borenstein et al 2010).

Maps of *p*-values resulting from the meta-analysis were corrected for multiple comparisons using a modified searchlight false discovery rate (FDR) procedure set to *p*<0.05 (for details on procedures and code, see Langers et al 2007). We applied this correction globally across all seven bilateral subcortical regions and measures (thickness, surface area) for each linear model, as well as locally (i.e., independently in each subcortical region and for each measure). Importantly, both correction methods provide opportunity for valid inferences in the sense of controlling for FDR. In the global case, we control the FDR over all regions simultaneously; in the local case, we interpret each region’s regression analysis for each measure independently of all other regions. See “Meta-analytic framework and correction for multiple comparisons” in the Supplement for more details.

## Results

We conducted analyses using a conservative global FDR-correction (i.e., correction across the surfaces of all structures assessed, see previous section). However, we also report local FDR-correction (i.e., correction across the surface for a single structure), which is more appropriate for examining effects within specific structures that we hypothesized would yield MDD-related effects based on previous independent investigations: hippocampus, amygdala, and striatum. Nevertheless, for comprehensiveness, we conducted and report below local FDR-corrected results from all seven bilateral subcortical structures.

### Global FDR-corrected effects

#### MDD versus CTL (and interaction effects with age and sex)

There were no significant differences between MDD and CTL, and no significant interactions between diagnostic group and age or sex.

#### EO versus CTL

Relative to CTL, EO had lower thickness in the hippocampus (Cohen’s *d* = −0.17) and amygdala (Cohen’s *d* = −0.16), and smaller surface area in the hippocampus (Cohen’s *d* = −0.18) and amygdala (Cohen’s *d* = −0.17). The strongest effects were primarily in the surface area of the SUB, CA2/3, and BLA. See Table 1 and Figure 1 for more details.

**Figure 1.**
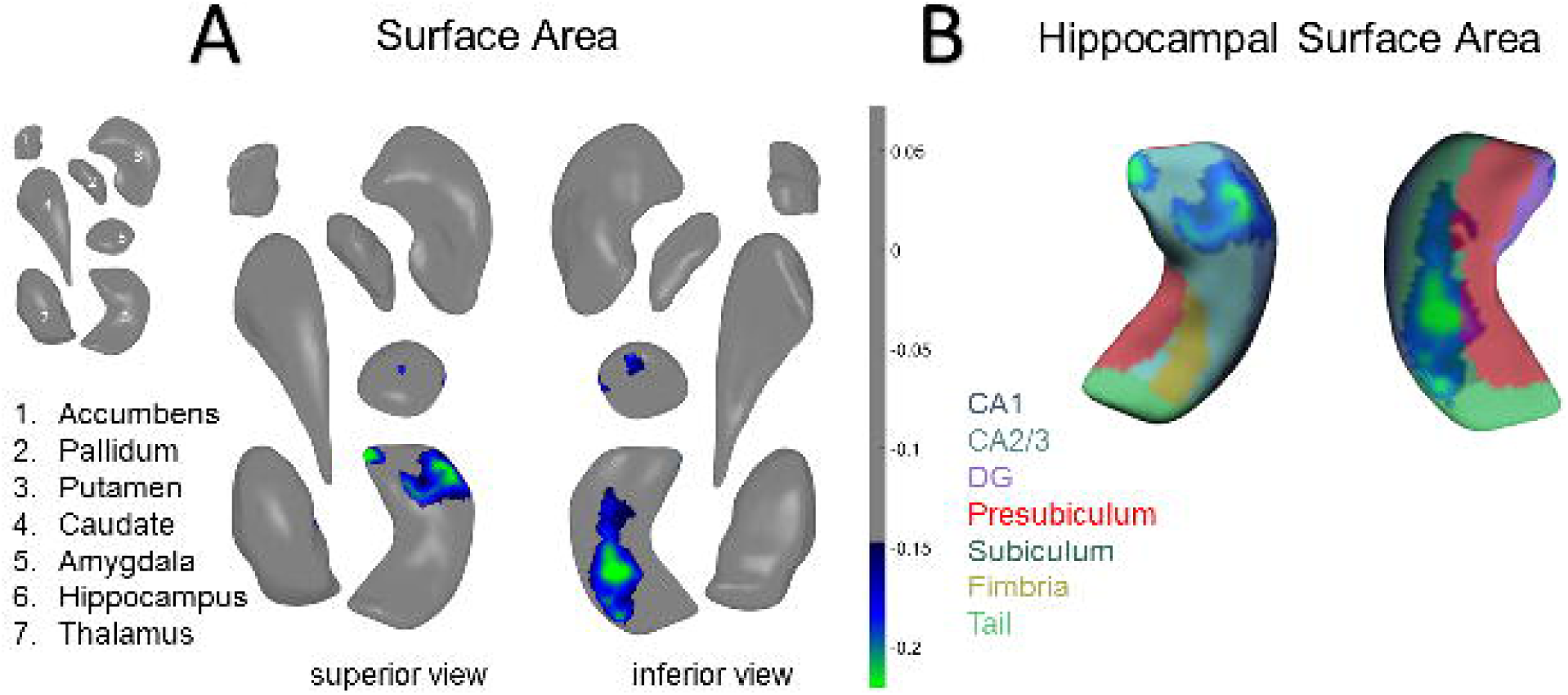
Global-FDR Corrected Results for EO v. CTL. **A)** Surface area effects in the amygdala and hippocampus from a superior view (left) and an inferior view (right) **B)** Surface area effects overlaid on the FreeSurfer v. 5.3 hippocampal subfield atlas (mirrored). Colored bars correspond to range of effect sizes (Cohen’s *d*). All results are based on bilateral shape measures (i.e., templates for corresponding left and right regions are vertex-wise registered after reflecting one of them, and summed vertex-wise). See Table 1 in the main text for more information.

#### LO versus CTL

There were no significant differences between LO and CTL.

#### EO versus LO

There were no significant differences between EO and LO.

#### RECUR versus CTL

There were no significant differences between RECUR and CTL.

#### FIRST versus CTL

There were no significant differences between FIRST and CTL.

#### RECUR versus FIRST

Relative to FIRST, RECUR had lower thickness in the hippocampus (Cohen’s *d* = −0.17) and amygdala (Cohen’s *d* = −0.17), and smaller surface area in the hippocampus (Cohen’s *d* = −0.17) and amygdala (Cohen’s *d* = −0.18). These effects were primarily in the surface area of the CA1 and BLA. Relative to FIRST, RECUR also had both greater thickness (Cohen’s *d* = 0.18) and greater surface area (Cohen’s *d* = 0.18) in the medial posterior thalamus. See Table 1 and Figure 2 for more details.

**Figure 2.**
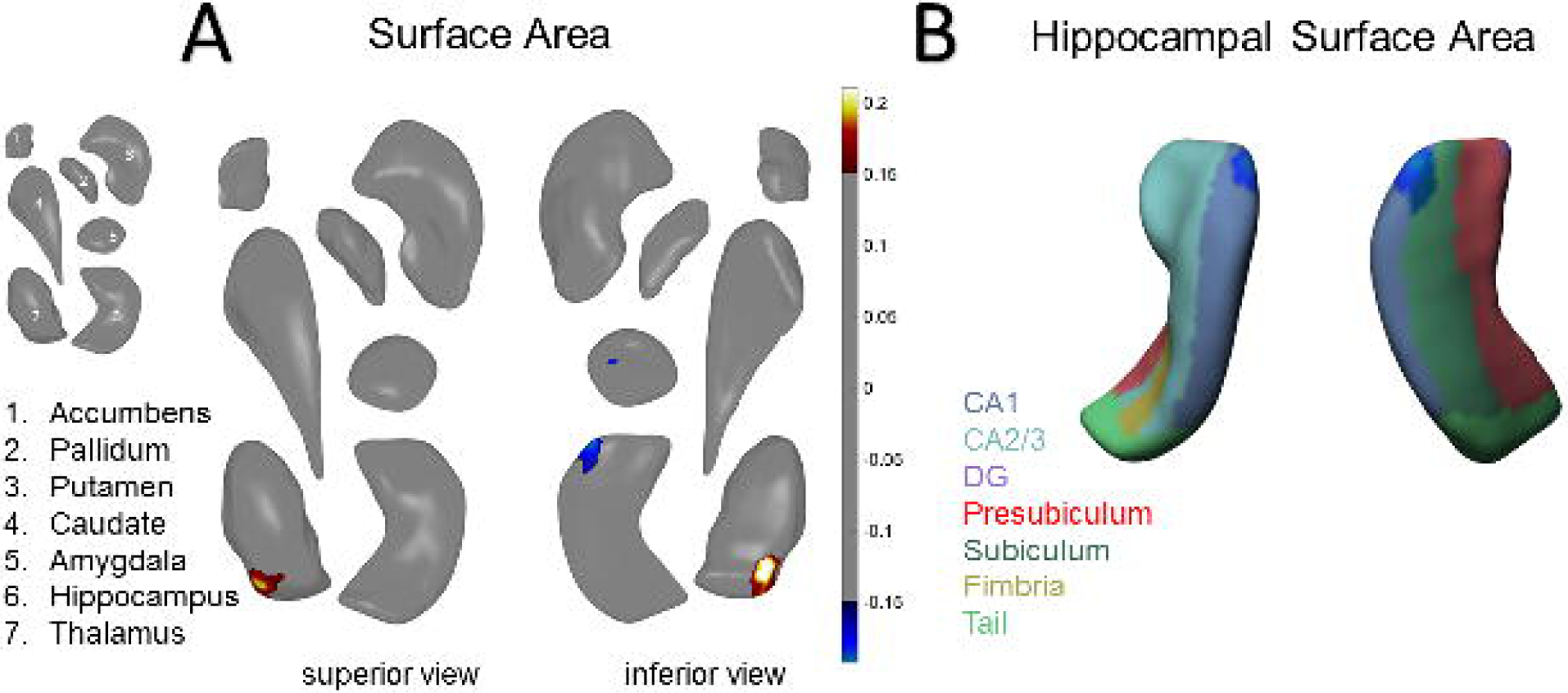
Global-FDR Corrected Results for RECUR v. FIRST. **A)** Surface area effects in the amygdala, hippocampus, and thalamus from a superior view (left) and an inferior view (right) **B)** Surface area effects overlaid on the FreeSurfer v. 5.3 hippocampal subfield atlas. Color bars correspond to range of effect sizes (Cohen’s *d*). All results are based on bilateral shape measures (i.e., templates for corresponding left and right regions are vertex-wise registered after reflecting one of them, and summed vertex-wise). See Table 1 in the main text for more information.

#### Associations with Depression Severity

There were no significant associations with depressive symptom severity using HDRS-17 or BDI scores in any subcortical structural outcome measures.

### Local FDR-corrected effects

#### MDD versus CTL (and interaction effects with age and sex)

Relative to CTL, MDD exhibited lower caudate thickness (Cohen’s *d* = −0.20) and smaller hippocampal (Cohen’s *d* = −0.11), amygdala (Cohen’s *d* = −0.11), and NAcc (Cohen’s *d* = −0.12) surface area. These effects were found for the surface area of SUB, BLA, and NAcc-s. See Table 1 and Figure 3 for more details. We found no significant interaction effects of diagnostic group with either age or sex.

**Figure 3.**
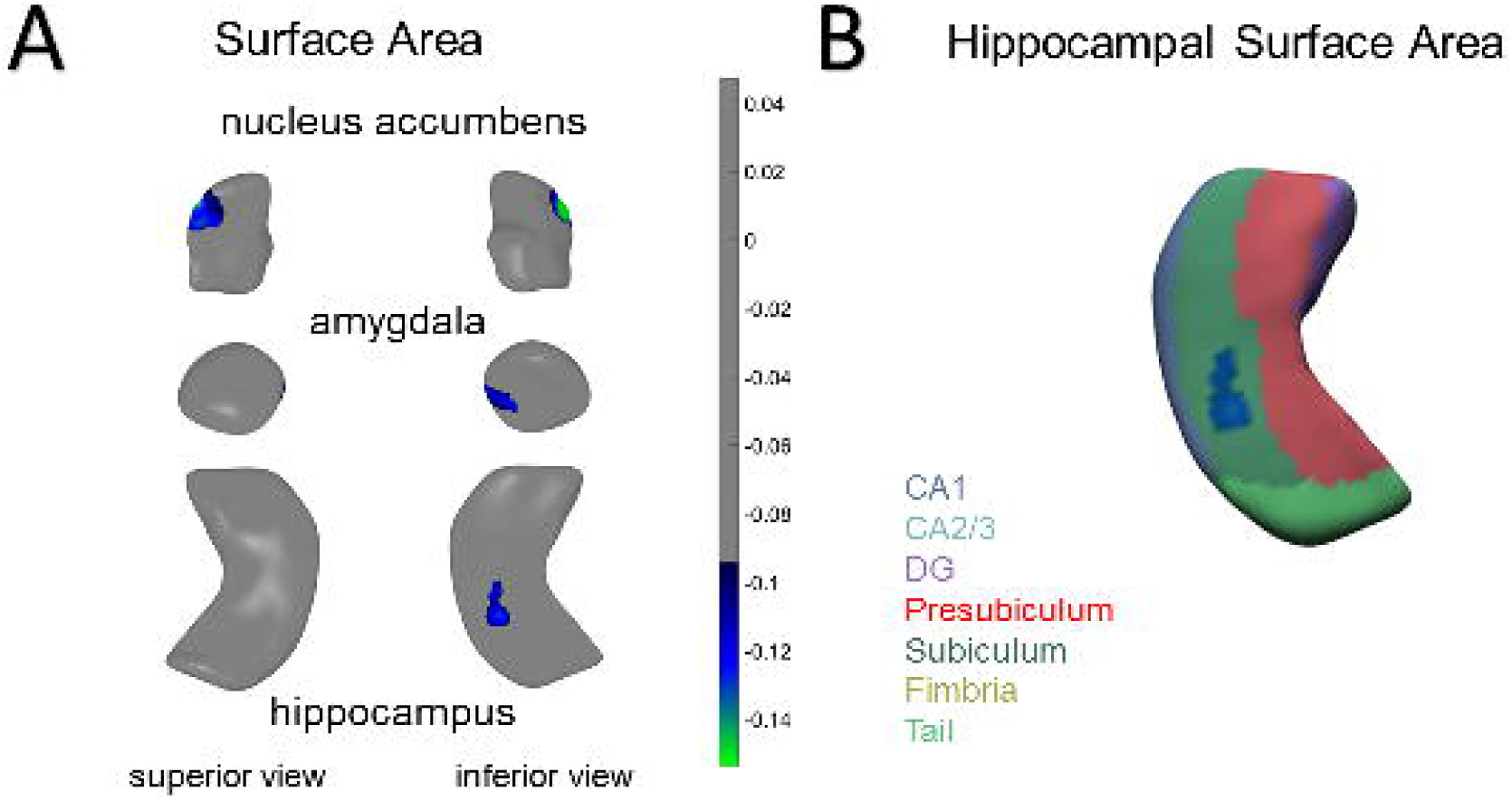
Local-FDR Corrected Results for MDD v. CTL. **A)** Surface area effects in the amygdala, hippocampus, and nucleus accumbens from an inferior view (right) **B)** Surface area effects overlaid on the FreeSurfer v. 5.3 hippocampal subfield atlas. Color bars correspond to range of effect sizes (Cohen’s *d*). All results are based on bilateral shape measures (i.e., templates for corresponding left and right regions are vertex-wise registered after reflecting one of them, and summed vertex-wise). See Table 1 in the main text for more information.

#### EO versus CTL

Relative to CTL, EO exhibited lower hippocampal (Cohen’s *d* = −0.17) thickness, and smaller hippocampal (Cohen’s *d* = −0.17) and amygdala (Cohen’s *d* = −0.17) surface area. These effects were found primarily in the SUB and BLA, consistent with the global FDR-corrected results. See Table 1 and Figure S1 for more details.

#### LO versus CTL

Relative to CTL, LO exhibited smaller nucleus accumbens (Cohen’s *d* = −0.13) and caudate (Cohen’s *d* = −0.17) surface area. These effects were found in the NAcc-s and caudate body. See Table 1 and Figure S1 for more details.

#### EO versus LO

There were no significant differences between EO and LO.

#### RECUR versus CTL

Relative to CTL, RECUR exhibited lower amygdala (Cohen’s *d* = −0.13) thickness, and smaller amygdala (Cohen’s *d* = −0.13) and NAcc (Cohen’s *d* = −0.13) surface area. These effects were found primarily in the BLA and NAcc-s. See Table 1 and Figure S2 for more details.

#### FIRST versus CTL

Relative to CTL, FIRST exhibited smaller hippocampal (Cohen’s *d* = −0.17) and caudate (Cohen’s *d* = −0.15) surface area. These effects were found primarily in the CA2/3 regions and caudate head. See Table 1 and Figure S3 for more details.

#### RECUR versus FIRST

Relative to FIRST, RECUR exhibited lower amygdala (Cohen’s *d* = −0.17) as well as greater thalamic thickness (Cohen’s *d* = 0.17), and enlarged thalamic surface area (Cohen’s *d* = 0.16). These effects were found primarily in BLA amygdala and the medial posterior thalamus. See Table 1 and Figure S4 for more details.

### Supplemental results of group comparisons based on antidepressant usage

Please see Table 1 for a summary of these results at both global and local FDR-corrected thresholds, and the Supplemental Information and Figures S5-S7 for more details.

## Discussion

The present study represents the largest investigation of subcortical shape in MDD to date. We identified reductions in the thickness and surface area of the subiculum (SUB) of the hippocampus and the basolateral amygdala (BLA) that appear to be driven by groups of patients with an adolescent age of onset (i.e., prior to age 21 years). Further, recurrence of depression (i.e., more than one episode of MDD), respectively. Our results address limitations in our current understanding of neural substrates of MDD, revealing group differences that are manifested in nuanced changes in subcortical morphometry. These patterns may offer insight into important clinical influences on the brain basis of MDD.

In our initial study from the ENIGMA-MDD Working Group, we found significant reductions in hippocampal volume that were primarily driven by patients who had an age of onset of depression prior to 21 years (and/or patients experiencing recurrent episodes of MDD; Schmaal et al 2016). Our present finding that smaller surface area of the SUB differs between early-onset MDD and CTL suggests that stress plays a key role in the development of MDD, consistent with broader theoretical literature (Hammen 2005). The SUB receives input from other subfields of the hippocampus (especially CA1), has reciprocal connections with the hypothalamic nuclei, and sends projections to several subcortical and cortical targets, making it a key structure that regulates the HPA axis (Lowry 2002; O’Mara 2005). Post mortem data further indicate that human SUB may contain a higher density of glucocorticoid binding sites than CA1–4 or even the DG (Kim et al 2015; Sarrieau et al 1986). In this context, our results are consistent with preclinical and clinical models of MDD that posit that environmental stressors trigger depressive episodes through stress-induced increases in glucocorticoid levels that, in turn, shrink dendrites and reduce the number of spines in the hippocampus, resulting in atrophy (Frodl et al 2008; McEwen et al 2015; Tata and Anderson 2010). Given evidence that the SUB is especially vulnerable to environmental input early in life such that there may be sensitive or critical windows of development in this structure (Teicher et al 2012), our findings are also consistent with the idea that the effects of early-onset MDD on hippocampal volume may be partially explained by exposure to stress, including childhood adversity, which is strongly associated with an earlier age of MDD onset (Kessler et al 2010; McLaughlin et al 2010). Future longitudinal studies are needed to examine whether reductions in the hippocampus, and specifically in the SUB, mediate links between childhood maltreatment and the development of (early-onset) MDD.

In our present study, we report that patients with recurrent MDD exhibited reduced basolateral amygdala (BLA), reduced caudate body, reduced shell of the NAcc (NAcc-s), and enlarged medial posterior thalamus. Our finding of reduced BLA in patients with recurrent MDD relative to those in their first episode – as well as lower BLA in patients with recurrent MDD relative to CTL (in regional analyses) – clarifies conflicting data in the extant literature on the effects of MDD on amygdala volume and is consistent with the role of the BLA in responding to threats and stressors in the environment (Terburg et al 2018). Indeed, previous studies have documented that age of onset, number of episodes, and antidepressant medication affect amygdala volume in people with MDD (Hamilton et al 2008; Kronenberg et al 2009; Rubinow et al 2016; Schmaal et al 2016; van Eijndhoven et al 2009). Interestingly, in our supplemental analyses examining patients who were medicated (at the time of scan) and also those who were not medicated versus CTL, we did not find evidence of enlarged amygdala volume, as was reported in a meta-analysis from a decade ago (Hamilton et al 2008). Given some overlap in sites and samples, it is not surprising our results are more aligned with our prior meta-analysis of aggregate subcortical volumes, where we reported a trend that individuals with MDD exhibit reduced amygdala volume compared to healthy controls (Schmaal et al 2016). Thus, the use of shape analysis in this well-powered meta-analysis allowed us to not only detect more nuanced effects of MDD on amygdala morphometry but also to discover that recurrence of MDD is an important clinical characteristic associated with this neural marker.

In our prior meta-analysis of aggregate subcortical volumes in MDD, we did not detect differences between MDD and CTL in striatal volumes (Schmaal et al 2016). In contrast, in the regional analyses of the present meta-analysis, we detected smaller caudate body and shell of the NAcc (NAcc-s) in MDD compared to CTL and smaller NAcc-s in recurrent MDD compared to CTL. The NAcc-s receives projections primarily from other limbic structures, including the hippocampus and amygdala, and is regarded by several researchers to comprise, in part, mesolimbic pathways (Deutch and Cameron 1992) and the extended amygdala (Alheid and Heimer 1988). Work in animals has demonstrated that, compared to the NAcc core, the NAcc-s contains higher concentrations of both dopamine and serotonin; these two divisions of the NAcc also responded differentially to pharmacological and environmental challenges, with haloperidol (an antipsychotic) affecting dopamine metabolism more in the core but stress (via an immobilization paradigm) selectively increasing dopamine release in the NAcc-s (Deutch and Cameron 1992; Scheggi et al 2002). Interestingly, injecting phencyclidine into the NAcc-s – but not the core – results in reward activity, suggesting that the NAcc-s is specifically implicated in reward effects (Carlezon and Wise 1996). Growing evidence indicates that neurons in the NAcc-s are involved in several processes that are disturbed in MDD, including encoding pleasure signals, integrating motivational salience, and supporting reward-based reinforcement learning (Heller et al 2009; Misaki et al 2016; Whitton et al 2015). While more research in this area is clearly needed, our results are consistent with neurobiological models of anhedonia and melancholic MDD that implicate mesolimbic dysfunction and suggest that the NAcc-s – along with the BLA – may be candidate treatment targets or biomarkers.

Finally, our finding of greater thalamic thickness and surface area in recurrent patients with MDD relative to first-episode patients is an intriguing result that requires more research. While one study of postmortem samples reported more neurons in the mediodorsal and anteromedial nuclei of the thalamus in people diagnosed with MDD relative to CTL (Young et al 2004), others have reported larger thalamic volumes in first-episode treatment-naïve patients with MDD (Qiu et al 2014; Zhao et al 2014). Interestingly, in a meta-analysis by Bora et al., late-life depression was associated with smaller thalamic volume (Bora et al 2012). Lithium usage is associated with larger thalamic volumes in patients with bipolar disorder (Lopez-Jaramillo et al 2017; Lyoo et al 2010), but it is unclear from our data as well as in the current literature what the role of mood stabilizing medications are on brain structure in patients with MDD. As we report in the Supplementary Information, patients receiving antidepressant treatment at the time of scan did not differ, on average, in thalamus thickness or surface area compared to CTL. It will be important for future research to carefully consider the role of the thalamus in MDD and determine how illness recurrence and/or medication usage affects morphometry of this structure.

It is worth noting some nuances in interpreting results from the two FDR-correction approaches in light of our global analyses indicating smaller CA1 in patients with recurrent depression versus first-episode depression but no differences in hippocampal shape between these two clinical groups in our regional analyses. Generally, local-FDR correction is more permissive: if some location in one of the regions passes the global-FDR threshold for significance, then some location in one of the regions will pass that region’s local threshold. However, these two regions may or may not be the same. For example, suppose there are two regions, one of which exhibits a greater overall effect than the other. That is, the local critical *p*-value will be higher (i.e., more permissive) for the first and lower (i.e., more stringent) for the second, while the global critical *p*-value will lie somewhere between the two localized values. Now, suppose some location in the first meets significance at the global threshold, but no location for the second region meets significance at its respective local threshold. In this instance, when viewing these results from the basis of a global-FDR correction only, the first region would appear to exhibit a more significant effect (even though it actually has a relatively smaller effect compared to the second region). Thus, both correction methods provide opportunity for valid inferences in the sense of controlling for FDR; we therefore tested and report results from both global-FDR and local-FDR correction thresholds.

Overall, our effect sizes are small; nevertheless, they are comparable to what we have reported in prior meta-analytic investigations comparing MDD and CTL in subcortical and cortical regions (Schmaal et al 2017; Schmaal et al 2016). Further, for several of our analyses, the percentage of surface area or thickness of the subcortical structure demonstrating significant effects was sizable; for instance, relative to controls, patients with early-onset MDD showed a reduction of over 22% of hippocampal surface area. Given the heterogeneity of MDD as a disorder and the likely clinical heterogeneity across the different study sites (e.g., illness duration, medication usage), it may be that several of the findings we report here represent nuanced yet core variations in subcortical subregions as a function of clinical characteristics in MDD (e.g., early-onset depression, recurrent depression, etc).

### Strengths, limitations, and future directions

As the first multi-site meta-analytic study of subcortical shape in MDD, major strengths of our investigation include the large number of observations sampled from several sites across the world combined with the use of harmonization and standardized quality control across all of these sites. Our large sample size provided us adequate statistical power to detect nuanced effects of MDD on subcortical shape and also allowed us to include and correct for all 7 subcortical structures in our analyses.

Despite the harmonized preprocessing protocols and statistical analyses, one limitation of our meta-analytic investigation is that because we combined pre-existing data across worldwide samples, data collection protocols (e.g., scan sequences, depression measurements) were not harmonized. Therefore, there may be important sources of heterogeneity in both imaging acquisition protocols and in clinical assessments that will need to be considered in future investigations. Finally, as we alluded to previously, investigating the effects of antidepressant medication was challenging in the present study, as the majority of sites did not collect detailed information on history, duration/adherence, type, and dosage of antidepressant treatment. Future research studies focused on collecting detailed information on lifetime, as well as current, medication usage in patients with MDD are needed, to better understand how various antidepressants affect brain structure.

## Conclusions

We identified reductions in stress-sensitive subfields of the hippocampus, particularly in the subiculum, and in the basolateral amygdala in patients MDD compared to CTL; these effects were driven by patients with an earlier onset of depression. Examining nuances in subcortical shape may help disentangle the complex clinical influences on the brain basis of MDD and potentially identify intervention more precise intervention targets or more sensitive biomarkers of treatment response.

## Acknowledgments & Funding

This work was supported by NIH grants U54 EB020403 to PMT and R01 MH116147 to PMT.

The Study of Health in Pomerania (SHIP) is part of the Community Medicine Research net (CMR) (http://www.medizin.uni-greifswald.de/icm) of the University Medicine Greifswald, which is supported by the German Federal State of Mecklenburg-West Pomerania. MRI scans in SHIP and SHIP-TREND have been supported by a joint grant from Siemens Healthineers, Erlangen, Germany and the Federal State of Mecklenburg-West Pomerania.

The FOR2107 cohort was supported by the German Research Foundation (DFG, grant FOR2107 DA1151/5-1 and DA1151/5-2 to UD; SFB-TRR58, Projects C09 and Z02 to UD; grant FOR2107 KR 3822/7-2 to AK; FOR2107 KI 588/14-2 to TK, FOR2107 NE 2254/1-2 to IN and FOR2107 JA 1890/7-2 to AJ), the Interdisciplinary Center for Clinical Research (IZKF) of the medical faculty of Münster (grant Dan3/012/17 to UD).

DIP-Groningen cohort was supported by the Gratama Foundation, the Netherlands (2012/35 to NG).

The CODE cohort was collected from studies funded by Lundbeck and the German Research Foundation (WA 1539/4-1, SCHN 1205/3-1, SCHR443/11-1).

The Magdeburg-Sexpect cohort was supported by the German Research Foundation (DFG-SFB779/TPA06).

LS is supported by a NHMRC Career Development Fellowship (1140764). TCH is supported by NIH grant K01 MH117442. NJ is supported by NIH grants R01 MH117601, R01 AG059874, and U54 EB020403. PMT is supported in part by NIH grants U54 EB020403, R01 MH116147, R56 AG058854, R01 MH111671 and P41 EB015922. HJG is supported in part by the German Research Foundation (DFG), the German Ministry of Education and Research (BMBF), the DAMP Foundation, Fresenius Medical Care, the EU Joint Programme Neurodegenerative Disorders (JPND) and the European Social Fund (ESF). IHG is supported in part by NIH grant R37 MH101495. PGS is supported in part by the German Research Foundation (DFG, SA 1358/2-1) and the Max Planck Institute of Psychiatry, Munich.

HJG has received travel grants and speakers honoraria from Fresenius Medical Care and Janssen Cilag. KS has consulted for Roche Pharmaceuticals and Servier Pharmaceuticals. PMT has received partial research support from Biogen, Inc. unrelated to the topic of this manuscript. All other authors declare no biomedical conflicts of interest.

The funding agencies played no role in the design and conduct of the study; collection, management, analysis, and interpretation of the data; and preparation, review, or approval of the manuscript.

