## Supplemental Information for "Subcortical Shape Alterations in Major Depressive Disorder: Findings from the ENIGMA Major Depressive Disorder Working Group"

**Image processing and analysis**

We estimated shape using the ENIGMA-Shape protocol (<http://enigma.usc.edu/ongoing/enigma-shape-analysis/>), for which test-retest reliability has been previously validated (Hibar et al 2015). Shapes were extracted using the FreeSurfer 5.3 parcellation, followed by a topological correction and mild smoothing based on the topology-preserving level set algorithm (Gutman et al 2015), thereby representing each region as a triangulated surface model. Each surface was then spherically inflated and registered to a region-specific template model by matching local geometric features; in other words, each individual was represented by a set of shapes with vertex-to-vertex correspondence to a standard template surface (Gutman et al 2015; Gutman et al 2012). In the present study, the template we used was the ENIGMA-Shape atlas, which was constructed by computing the Euclidean average of the surface models of 200 (100 male) unrelated individuals from the Queensland Twin Imaging Study (QTIM); QTIM was not used in this study beyond atlas construction.

Following this registration procedure, we subjected each individual surface to a medial curve and further refined the registration as needed (Gutman et al 2015; Gutman et al 2012). We then defined two vertex-wise measures of space morphometry which facilitated comparisons of subcortical shape: *radial distance*, as derived from the medial model; (Gutman et al 2015; Gutman et al 2012), which yields “shape thickness,” and the *Jacobian determinant,* as derived from tensor based morphometry, TBM (Gutman et al 2015; Wang et al), which yields a metric of localized tissue reduction or enlargement of surface area (relative to the respective template shape). Because the Jacobian represents the ratio of the area in the individual shape relative to the area in the template at the corresponding vertex and not Gaussian in distribution, we used the logarithm of the Jacobian in all analyses examining shape surface area. A useful feature of the ENIGMA-Shape pipeline is that results are based on bilateral shape measures (i.e., templates for corresponding left and right regions are vertex-wise registered after reflecting one of them and summed vertex-wise). Importantly, our registration algorithm provides a unique and stable matching between datasets, allowing us to efficiently meta-analyze the effects of MDD across datasets (as in (Roshchupkin et al 2016)).

Visual quality control in all regions of interest was performed by individual raters according to the ENIGMA-Shape Quality Control guide (<http://enigma.usc.edu/ongoing/enigma-shape-analysis/>). Beyond the guide, sites performing local quality control were given access to experienced raters at the Imaging Genetics Center for particularly difficult cases.

**Meta-analytic framework and correction for multiple comparisons**

Maps of *p*-values resulting from the meta-analysis were corrected for multiple comparisons using a modified searchlight false discovery rate (FDR) procedure set to *p*<0.05 (for details on procedures and code, see Langers et al 2007). We applied this correction globally across all 7 bilateral subcortical regions and measures (thickness, surface area) for each linear model, as well as locally (i.e., independently in each subcortical region and for each measure). It is worth noting that both correction methods provide opportunity for valid inferences in the sense of controlling for FDR. In the global case, we control the FDR over all regions simultaneously; in the local case, we interpret each region’s regression analysis for each measure independently of all other regions.

In both cases, a Gaussian weighting function with 6 mm full-width half maximum (FWHM) kernel size was used for the searchlight with 2 mm Gaussian smoothing (Langers et al 2007). In the case of global correction, we assumed no correlation in *p*-values between vertices of different neighboring regions (i.e., by setting the Euclidean distance between vertices of different subcortical regions to infinity), resulting in a more conservative critical *p* estimate.

**Supplemental results of group comparisons based on antidepressant usage**

***Global-FDR correction***

*MED versus CTL*
Compared to CTL, MED exhibited reduced hippocampal (Cohen’s *d* = -0.139) and caudate (Cohen’s *d* = -0.133) thickness, and reduced hippocampal (Cohen’s *d* = -0.136), caudate (Cohen’s *d* = -0.140), and NAcc (Cohen’s *d* = -0.142) surface area. See Table 1 in the main text and Figure S6 for more details.

*NON versus CTL*
We found no significant differences between NON and CTL.

***Local-FDR correction***
 *MED versus CTL*
Compared to CTL, MED exhibited reduced caudate (Cohen’s *d* = -0.12) thickness, and reduced hippocampal (Cohen’s *d* = -0.13) and NAcc (Cohen’s *d* = -0.14) surface area. These effects were similar to results from the global FDR-corrected analyses. See Table 1 in the main text for more details.

*NON versus CTL*
Compared to CTL, NON exhibited reduced amygdala surface area (Cohen’s *d* = -0.14). See Table 1 in the main text and Figure S7 for more details.

**Table S1. Uploaded as a separate document.**

**Figure S1. Local-FDR Corrected Results for LO v. CTL.** Surface area effects in the nucleus accumbens and caudate from a superior view (left) and an inferior view (right). Color bars correspond to range of effect sizes (Cohen’s *d*). All results are based on bilateral shape measures (i.e., templates for corresponding left and right regions are vertex-wise registered after reflecting one of them, and summed vertex-wise). See Table 1 in the main text for more information.

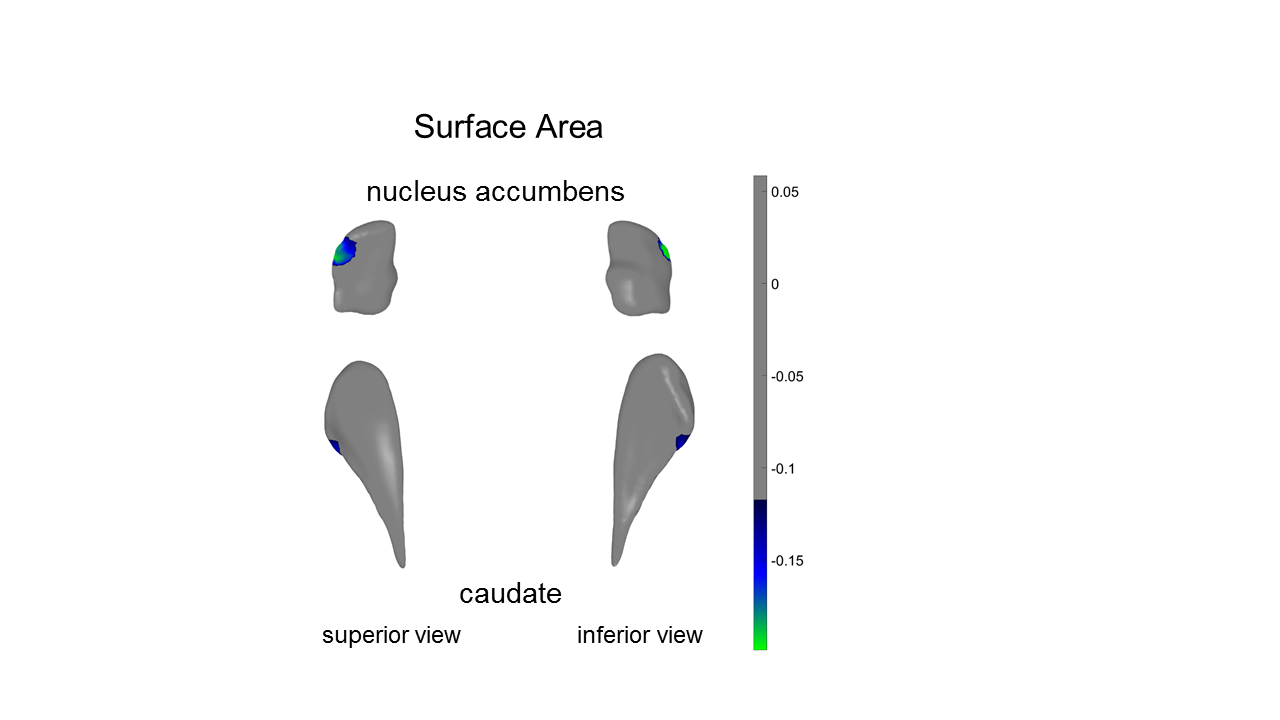

**Figure S2. Local-FDR Corrected Results for RECUR v. CTL.** **A)** Thickness effects in the amygdala and nucleus accumbens from a superior view (left) and an inferior view (right) **B)** Surface area effects in the amygdala and nucleus accumbens from a superior view (left) and an inferior view (right). Color bars correspond to range of effect sizes (Cohen’s *d*). All results are based on bilateral shape measures (i.e., templates for corresponding left and right regions are vertex-wise registered after reflecting one of them, and summed vertex-wise). See Table 1 in the main text for more information.

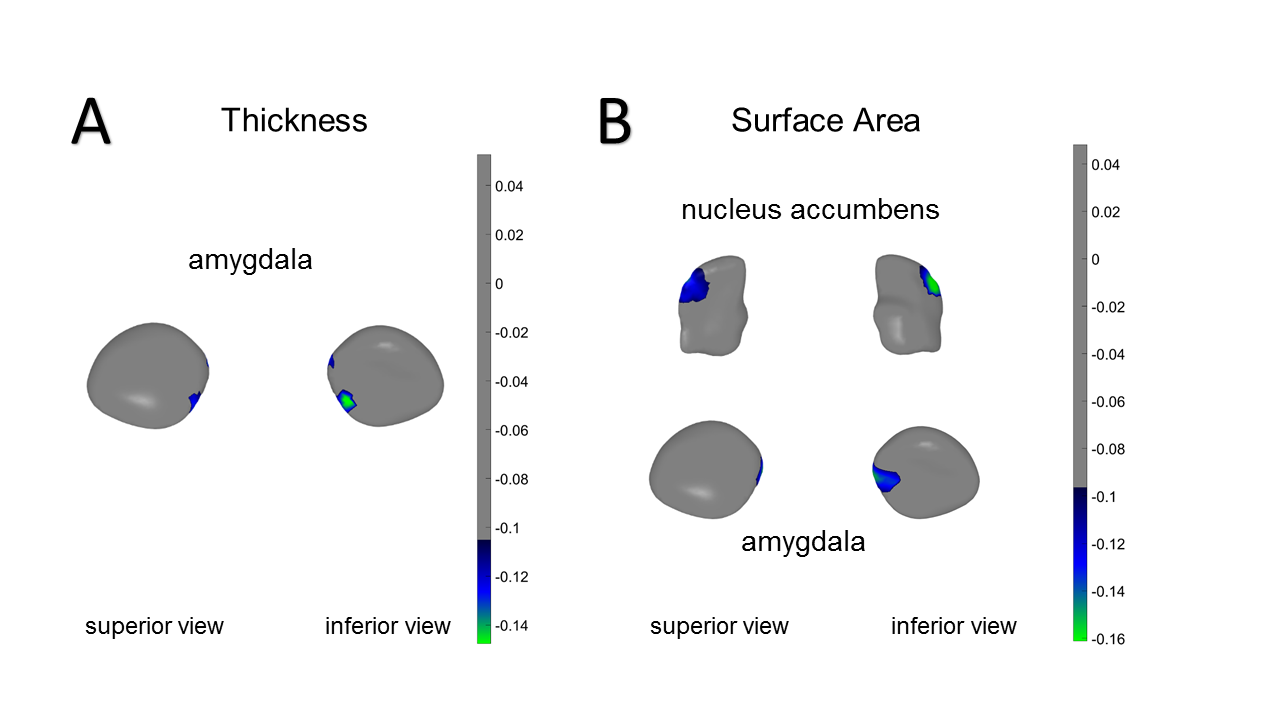

**Figure S3. Local-FDR Corrected Results for FIRST v. CTL.** Surface area effects in the hippocampus and caudate from a superior view (left) and an inferior view (right). Color bars correspond to range of effect sizes (Cohen’s *d*). All results are based on bilateral shape measures (i.e., templates for corresponding left and right regions are vertex-wise registered after reflecting one of them, and summed vertex-wise). See Table 1 in the main text for more information.

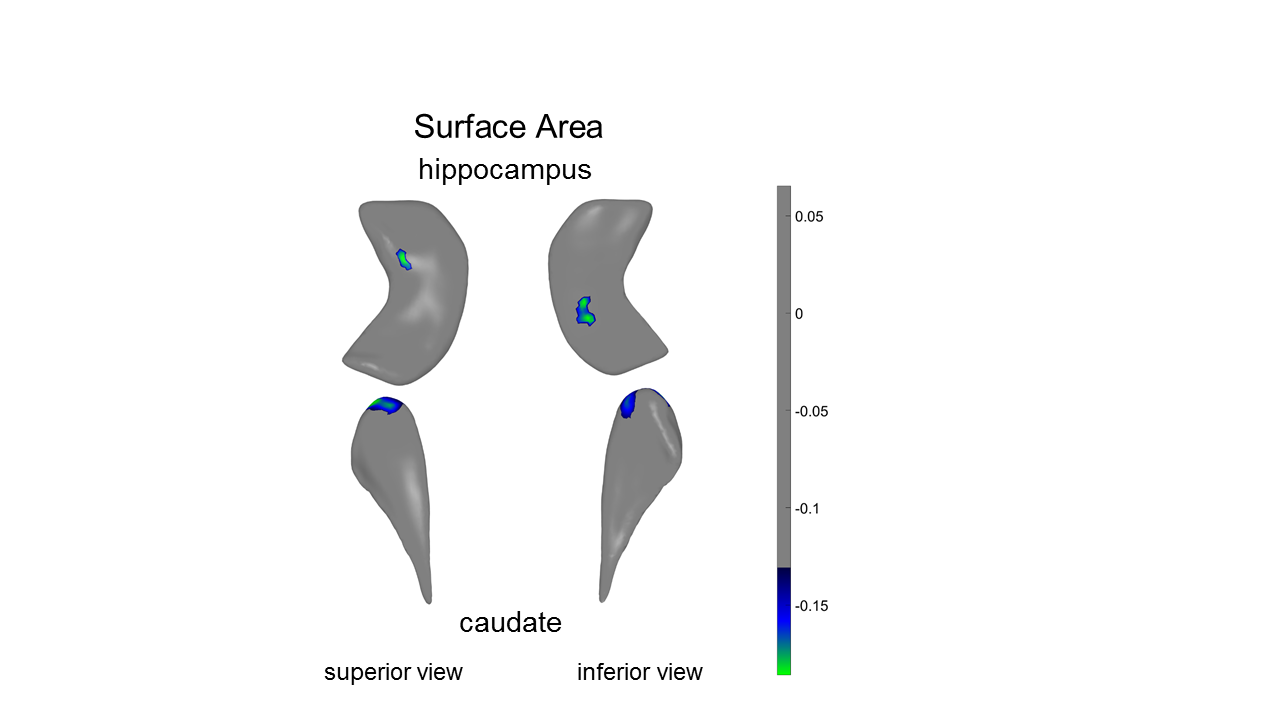

**Figure S4. Local-FDR Corrected Results for RECUR v. FIRST.** **A)** Thickness effects in the amygdala and thalamus from a superior view. **B)** Surface area effects in the thalamus shown from a superior view. Color bars correspond to range of effect sizes (Cohen’s *d*). All results are based on bilateral shape measures (i.e., templates for corresponding left and right regions are vertex-wise registered after reflecting one of them, and summed vertex-wise). See Table 1 in the main text for more information.

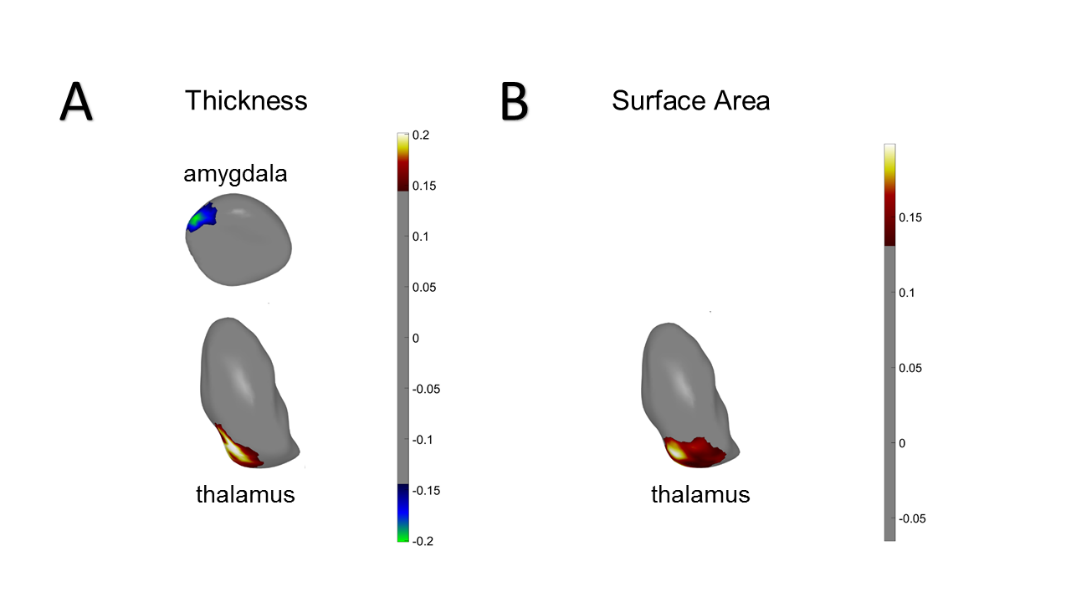

**Figure S5. Global-FDR Corrected Results for MED v. CTL.** **A)** Thickness effects in the hippocampus and caudate from a superior view (left) and an inferior view (right) **B)** Surface area effects in the hippocampus, caudate, and NAcc shown from a superior view (left) and an inferior view (right). Color bars correspond to range of effect sizes (Cohen’s *d*). All results are based on bilateral shape measures (i.e., templates for corresponding left and right regions are vertex-wise registered after reflecting one of them, and summed vertex-wise). See Table 1 in the main text for more information.

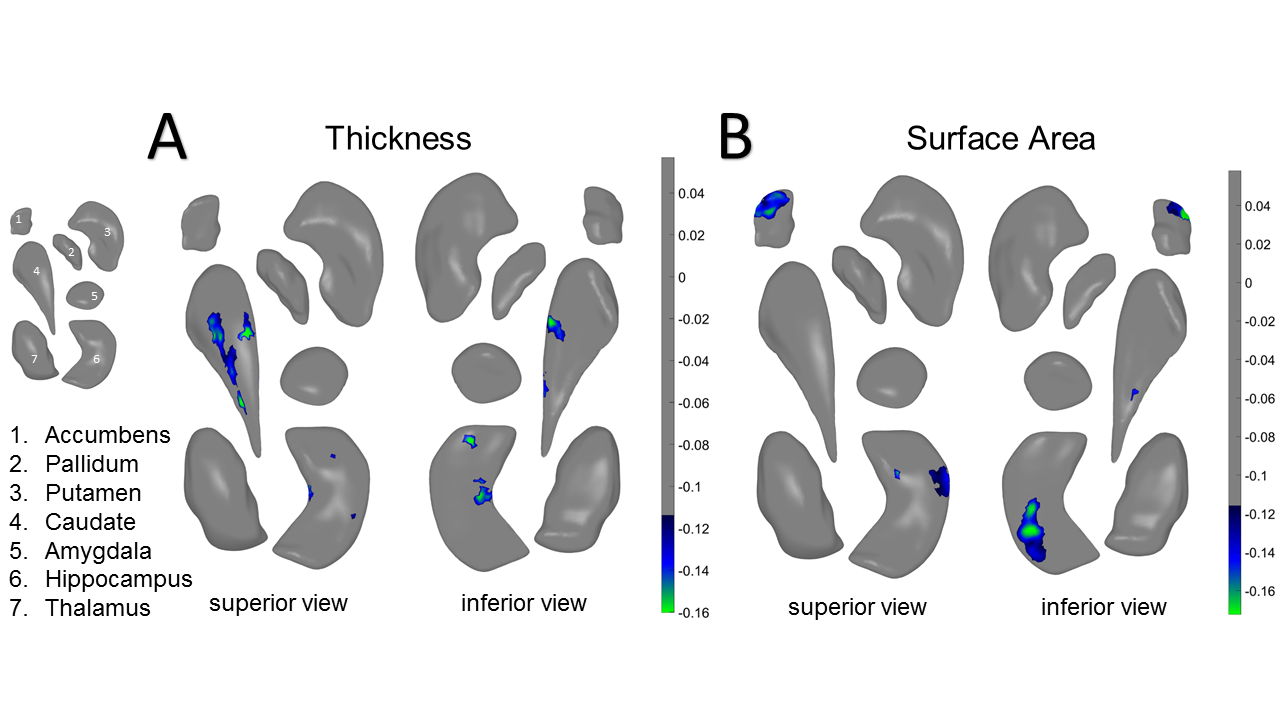

**Figure S6. Local-FDR Corrected Results for MED v. CTL.** **A)** Thickness effects in the caudate from a superior view (left) and an inferior view (right) **B)** Surface area effects in the hippocampus and NAcc shown from a superior view (left) and an inferior view (right). Color bars correspond to range of effect sizes (Cohen’s *d*). All results are based on bilateral shape measures (i.e., templates for corresponding left and right regions are vertex-wise registered after reflecting one of them, and summed vertex-wise). See Table 1 in the main text for more information.

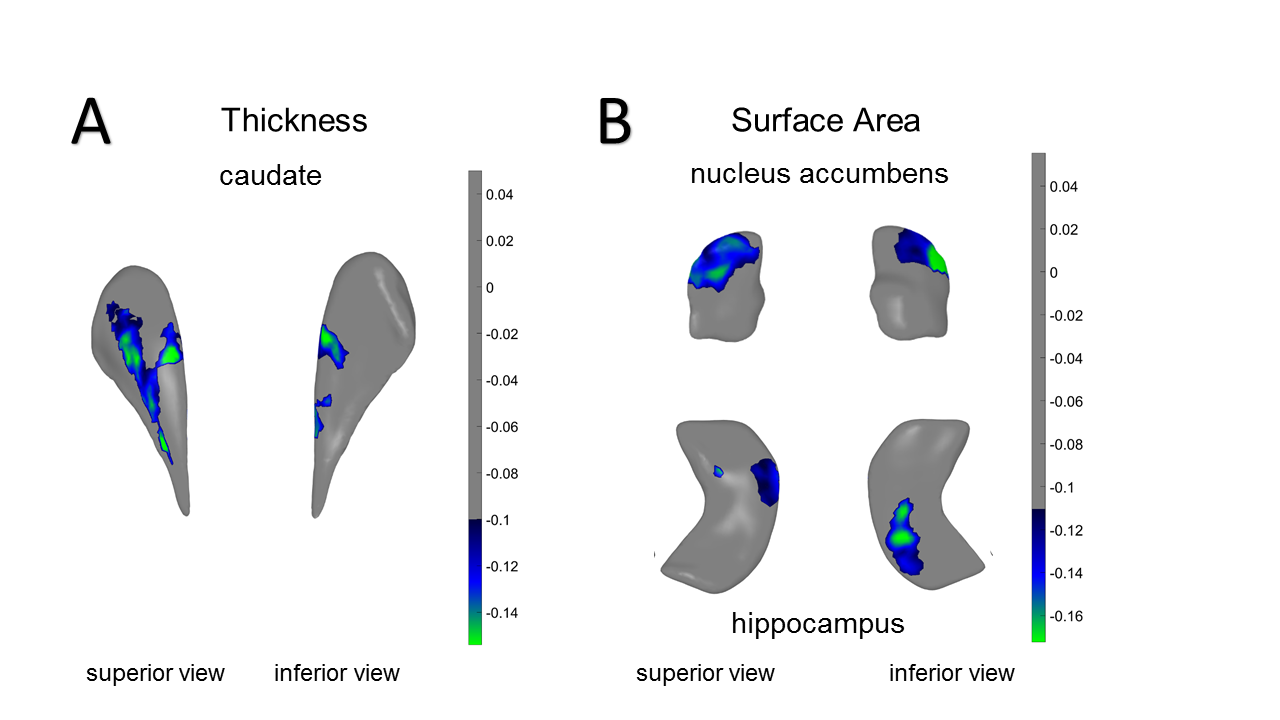

**Figure S7. Local-FDR Corrected Results for NON v. CTL.** Surface area effects in the amygdala shown from a superior view (left) and an inferior view (right). Color bars correspond to range of effect sizes (Cohen’s *d*). All results are based on bilateral shape measures (i.e., templates for corresponding left and right regions are vertex-wise registered after reflecting one of them, and summed vertex-wise). See Table 1 in the main text for more information.

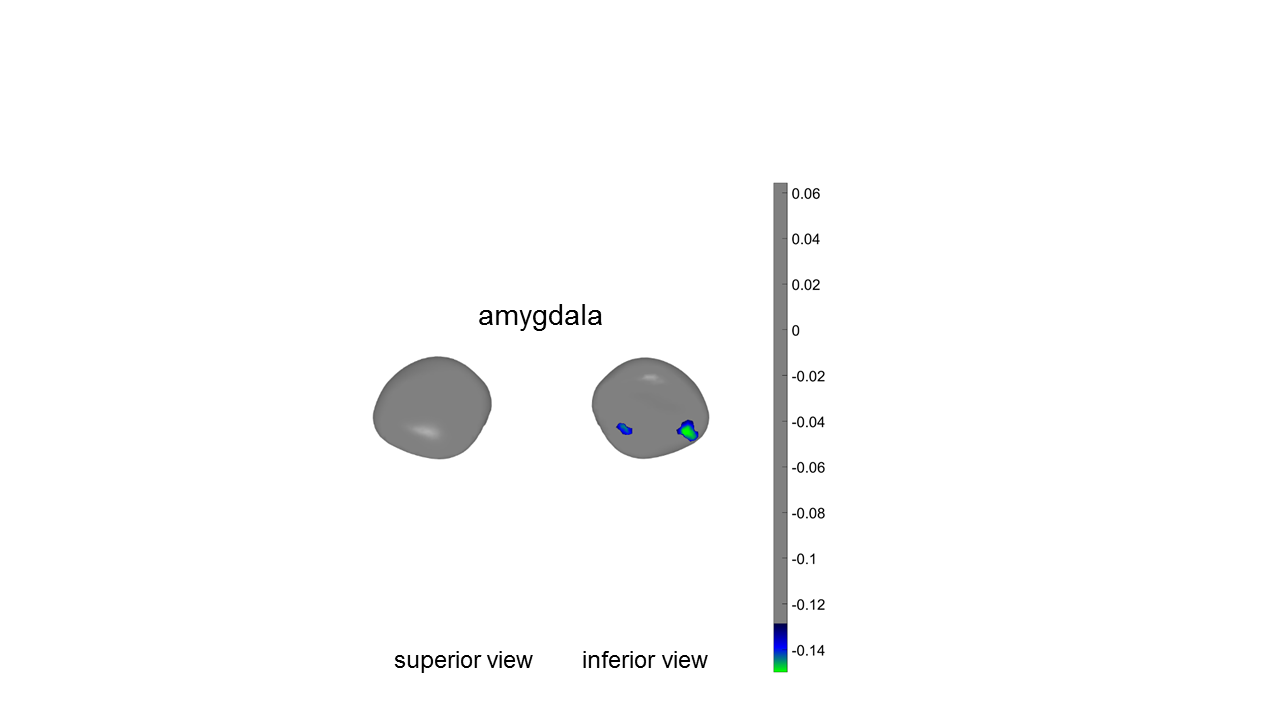
